# Popformer: Learning general signatures of positive selection with a self-supervised transformer

**DOI:** 10.64898/2026.03.06.710163

**Authors:** Leon Zong, Sorelle A. Friedler, Sara Mathieson

## Abstract

Understanding natural selection can help shed light on the genetics underpinning adaptive evolution. The widespread availability of large-scale human genetic variation data has led to the development of data-driven methods for detecting signatures of selection, many of which are based on deep learning. However, these methods often fail to generalize well to the diversity of selection signatures across a broad range of evolutionary scenarios. We propose a novel transformer-based model, Popformer, for learning encodings of general patterns of genetic variation. Popformer includes site-wise and haplotype-wise attention, allowing us to capture variation among both genetic positions and individuals. It additionally learns relative positional embeddings for inter-SNP distances. The model is pre-trained with an analog of the masked language modeling objective across a range of real human genomic data, similar to the task of genetic imputation. Using dimensionality reduction, we show that the pre-trained model learns meaningful embeddings of genomic windows that correspond with population structure. We also show that the model can accurately perform genotype imputation. We further fine-tune the model on selection classification and demonstrate that our model is more accurate than other selection classification methods, on selection simulations of both well-specified and mis-specified demographic models. Using a novel real data validation approach, we apply Popformer to human data from the 1000 Genomes Project and reveal its ability to generalize from simulated data to the diversity of real data. Overall, our method provides a new direction for population genetic inference methods, with future fine-tuning applications including the inference of recombination rates, introgression, and local ancestry.

**Author summary:** Natural selection lies at the heart of adaptation and evolution. Identifying genomic signals of natural selection can lead to insights about evolutionary pressures acting on populations, and provide an understanding of how populations differentiate. Detecting these signals using only present-day genomic data is difficult, and methods developed for this task often use specialized simulations, trading generalization for power. We show here that adapting deep learning techniques from modern language and image processing can be used to identify signals of selection with both high power and robustness to misspecification. By pre-training our transformer model, Popformer, on real data, we induce learned model representations that are representative of the variation in real data. We fine-tune our model on simulations inferred from European populations, and compare with other machine learning methods and summary statistics. Our results suggest that both our model architecture and our training regime are useful for strong performance in diverse simulation tests and in recovering known selection signatures in real data.

## Introduction

Modern population genetics broadly aims to understand patterns of variation in and between populations at a genome-wide level. A major goal of the field is to identify regions of the genome that are or previously were under natural selection. A beneficial mutation can rapidly rise in frequency in a population in a process known as a selective sweep. While this process leaves behind distinct patterns of variation in the population due to linked selection, identifying these sweeps is challenging [1, 2]. There are a number of confounding evolutionary effects, including those of demographic events and other stochastic forces like background selection and varying mutation and recombination rates [3–5]. Traditionally, theoretically motivated summary statistics are used to identify genetic regions which have undergone a selective sweep. These summary statistics identify a particular signature of selection; for example, reduced diversity (*π*, Tajima’s D), long stretches of identical DNA (iHS), or are a composite of several of these statistics (SweepFinder, CMS) [6–10]. While these methods are widely used, they are often underpowered and unreliable when confounding evolutionary forces produce the same genetic signatures [4, 11–13]. A new cohort of machine-learning based methods, many of which utilize convolutional neural networks (CNNs), have made progress in detecting selective sweeps with increased power and in the face of confounding evolution [14–21].

These new sweep detection methods have arisen in part due to the development of improved population genetics simulation programs including `SLiM`, `discoal`, and `msms`, which have the ability to model selective events [22–24]. With selection as a simulatable parameter, the methods are able to use a supervised training regime. Machine learning detection methods have proven to be powerful at detecting selection in well-specified matched simulations, and many have even demonstrated robustness to simulation misspecification [25, 26]. While success in the simulated regime is an important step, the ultimate goal of these methods is to discover signatures of selection in real genomic data. It is yet unclear to what extent models trained on simulations can generalize to real data. Simulations model theories of evolutionary processes, which are ultimately simplifications of the complex nature of real evolution.

Here, we introduce Popformer, a transformer-based model which learns general signatures of variation from real population genetic data. The model operates on sets of variants in the form of haplotype matrices. By training on real variants, the model is able to learn common patterns of variation that are derived from real evolutionary processes. The model is pre-trained with a version of the self-supervised masked language modeling objective, in which random positions in the input matrices are masked and the model learns to recover the value at the masked positions [27]. We demonstrate that the pre-trained model can perform a limited form of genotype imputation on par with widely used methods. We additionally show that pre-training produces model embeddings which are good representations of genetic variation.

The pre-trained model could further be trained (fine-tuned) for region-level tasks such as predicting signatures of selection, recombination rate, or mutation rate. In this work, our downstream focus is primarily selection detection. We introduce new, diverse simulated selection datasets for training and testing supervised selection detection methods. On these datasets, we demonstrate that our method can detect selection with increased power compared to both traditional summary statistics as well as recent deep learning methods trained on the same data. We find that Popformer can generalize better than other methods at detecting selection in out-of-distribution simulated test scenarios. Additionally, we use reported lists of real selected sites [28, 29] as validation, and find that our model predicts enriched scores for these reported sites over baseline. As such, a trained Popformer model can be used out-of-the-box to perform selection detection on a wide range of populations without the need for per-dataset simulations and training.

The architecture of the model additionally lends itself to SNP-level and haplotype-level prediction tasks, such as local ancestry inference or introgression detection. While we do not explore this in this work, we hope to facilitate these directions by releasing checkpoints for the pre-trained Popformer-base model, fine-tuned selection classification model, simulated datasets, along with all code used for training and evaluation. All resources can be found at https://popformer.leonzong.com.

## Results

### Overview of Popformer method

Popformer is an encoder-only transformer model which takes haplotype matrices as input and outputs a vector representation (embedding) of each position of the matrix (further details can be found in the Methods). The encoder uses an implementation of axial attention: where attention is computed both along the variants for each haplotype, and along the haplotypes for each variant (Fig 1) [30, 31]. This allows the model to dynamically weight particular haplotype variants based on the full population and window context. Distances between variants are included as a learned positional attention bias, allowing the model to learn patterns of variant density.

**Fig 1.**
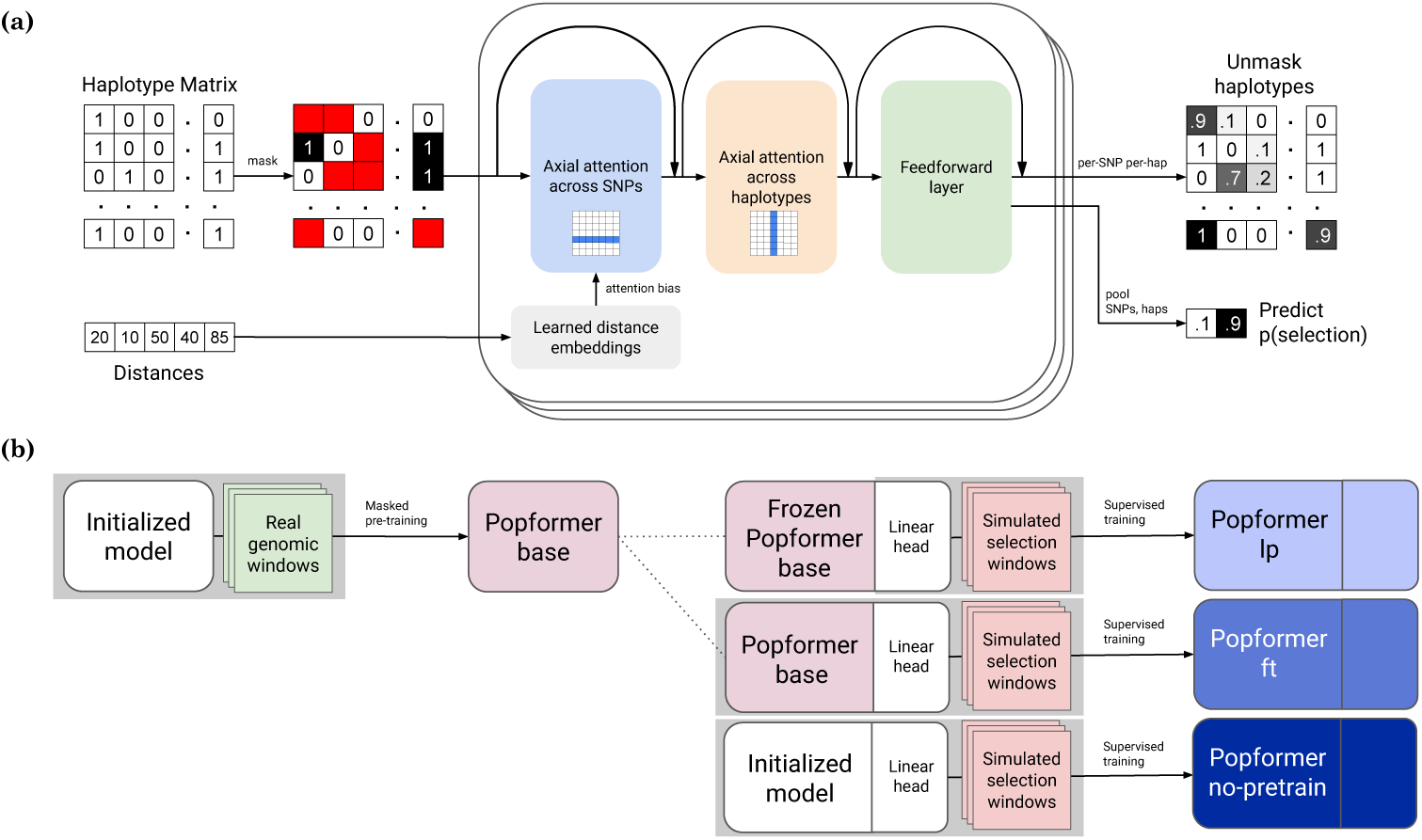
**(a)** Overview of the architecture and pre-training strategy of Popformer. Haplotype matrices and distance vectors are input to the model. The model consists of multiple blocks of axial attention. Each block consecutively applies SNP-wise attention across every haplotype, haplotype-wise attention for every SNP, and a fully-connected feed-forward layer. Normalized residual connections are applied after each layer. Pre-training is self-supervised, using real genomic windows. Random tokens are replaced with a mask token, and the model is tasked with unmasking the token. **(b)** Popformer-base is first trained using self-supervised unmasking. Using it, we evaluate three different selection training recipes. For Popformer-lp, the pre-trained base model is frozen and the head is trained on the static embeddings. For the Popformer-ft, gradients from training also flow through the base model. For Popformer-no-pretrain, we ablate the pre-training step, starting from a newly initialized model. The gray background indicates where gradients are computed.

During pre-training, the embedding representation is learned by training the model to unmask random positions in matrices derived from real human populations in the 1000 Genomes dataset [32]. We refer to this model as Popformer-base. On the downstream task of selection detection, we simulate a set of diverse genomic regions under human-inferred demographies, including both neutrally evolving regions and those evolving under various selection strengths. We use these simulations to fine-tune a simple linear model (the classification head) on top of Popformer-base under several different training recipes. We evaluate:

1. A linear probe (Popformer-lp), where the base model (encoder) is frozen and only the linear head is allowed to update.
2. Full fine-tuning (Popformer-ft), where both the base model and the linear head are allowed to update.
3. A pre-training ablation (Popformer-no-pretrain), where a newly initialized encoder is used in place of the pre-trained base model. Here, both the encoder and the head are allowed to update.

Once trained, these models are verified on held-out test simulations, where the ground truth is known. Additionally, they can be used to perform windowed inference along real genomes.

### Learned population embeddings capture informative genomic variation

Popformer-base is trained with an analogue of the masked language modeling task (see Methods) [27]. In this modeling scheme, the input sequences to be masked are haplotype matrices (S1 Fig). In order to ensure that our model accurately learns complex dependencies between SNPs, we first test the actual performance of our method on its training objective by computing the unmasking accuracy on a test set of 75%-uniformly-masked samples from 1000 Genomes populations. On this test, Popformer-base outperforms two simple baselines and performs quite well. The nearest neighbor baseline simply chooses the allele using the most similar windowed haplotype to the masked haplotype, and the allele frequency baseline chooses the most frequent unmasked allele (Table 1).

**Table 1.** Mean denoising accuracy and standard deviation for Popformer-base and two naive baselines. 75% of positions are uniformly masked from haplotype matrices of size 128 SNPs by 128 haplotypes, as in S1 Fig.

| Method | Accuracy |
| --- | --- |
| column frequency baseline | $90.3 \pm 2.96$ |
| nearest neighbor baseline | $88.3 \pm 2.49$ |
| **popformer-base** | **$95.8 \pm 1.17$** |

The task of unmasking haplotypes is also conceptually similar to the task of genetic imputation, whereby a ‘target’ panel of individuals has missing variant data imputed using a ‘reference’ panel. Our masked training differs from genotype imputation primarily in the nature of masking - in our case, we did not hold out a set of unmasked haplotypes to use as ‘reference’. Instead, all haplotypes were randomly masked. To compare to methods specifically designed for genotype imputation, we formulated an imputation-like test to further validate our pre-training objective performance (S2 Fig). *R*^2^ for genotype dosage imputation is calculated by first converting haplotypes into allele dosages (0, 1, or 2 for diploids) and then computing the squared Pearson correlation. Comparing *R*^2^, Popformer-base performs as well as IMPUTE5 [33], a state-of-the-art HMM imputation method, and continues to outperform both baselines (Fig 2). However, our method is worse at imputing exactly correct genotypes, consistently having a higher error rate. These results are quite promising given that our self-supervised training data has no special examples with a reference-vs-target masking pattern, and demonstrate that the pre-training strategy effectively learns patterns of genomic variation.

**Fig 2.**
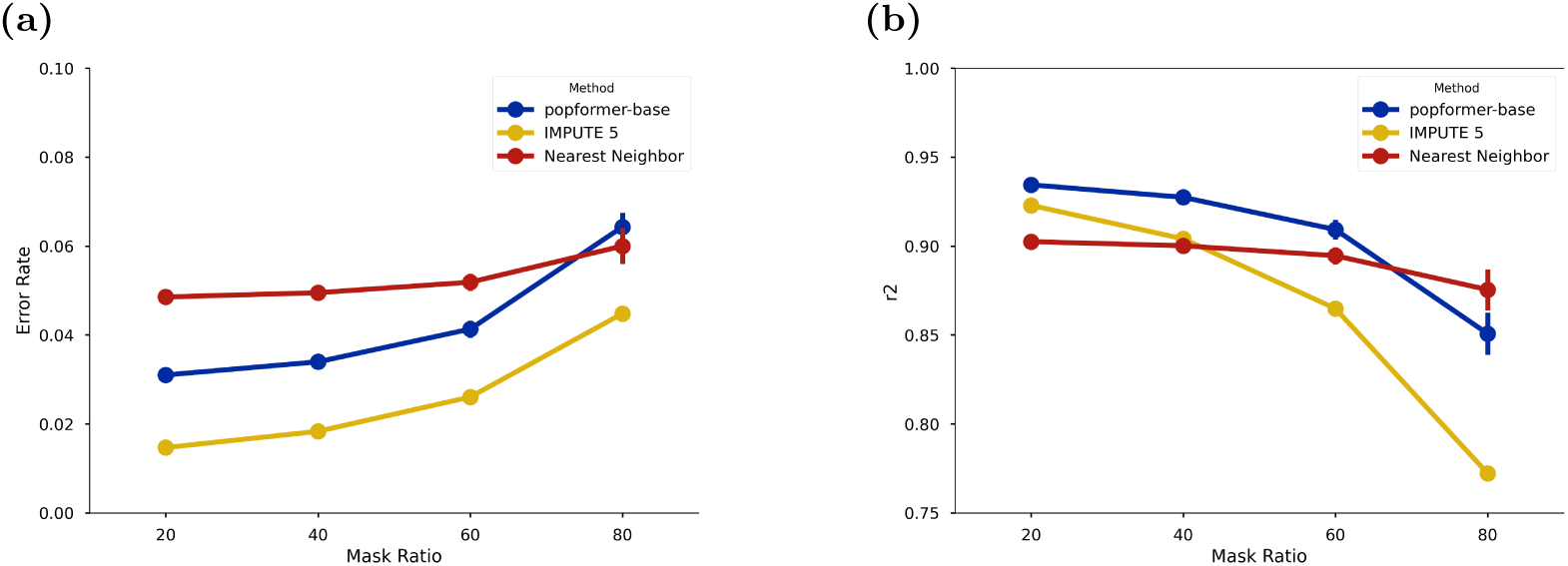
**(a)** Imputation error rate (lower is better) and **(b)** *R*^2^ (higher is better) at four different imputation levels. 64 haplotypes are chosen as an unmasked reference panel. 64 other haplotypes are used as a target panel, where a percentage of SNPs are randomly chosen to be masked. We compare performance for Popformer, IMPUTE5, and the nearest neighbor baseline. The column frequency baseline is omitted, performing significantly worse than the other methods.

While trained on unmasking, Popformer is primarily designed for applications in downstream tasks. In order to achieve good performance on these tasks, it is important that the model’s pre-training objective has produced an informative learned embedding. To inspect these representations, we first examined whether the embeddings adequately represent real genomic variation between populations at least at a continent-level scale. Individual embeddings were calculated for each individual in the 1000 Genomes dataset by concatenating mean-pooled window-level embeddings for sliding windows across chromosome 1 [32].

We performed a principal component analysis (PCA) on the these mean-pooled embeddings and used the first two components to visualize them (Fig 3a). Popformer-base’s embeddings match commonly reported results in which dimensionality-reduced genomic variation differentiates continental populations [34, 35].

**Fig 3.**
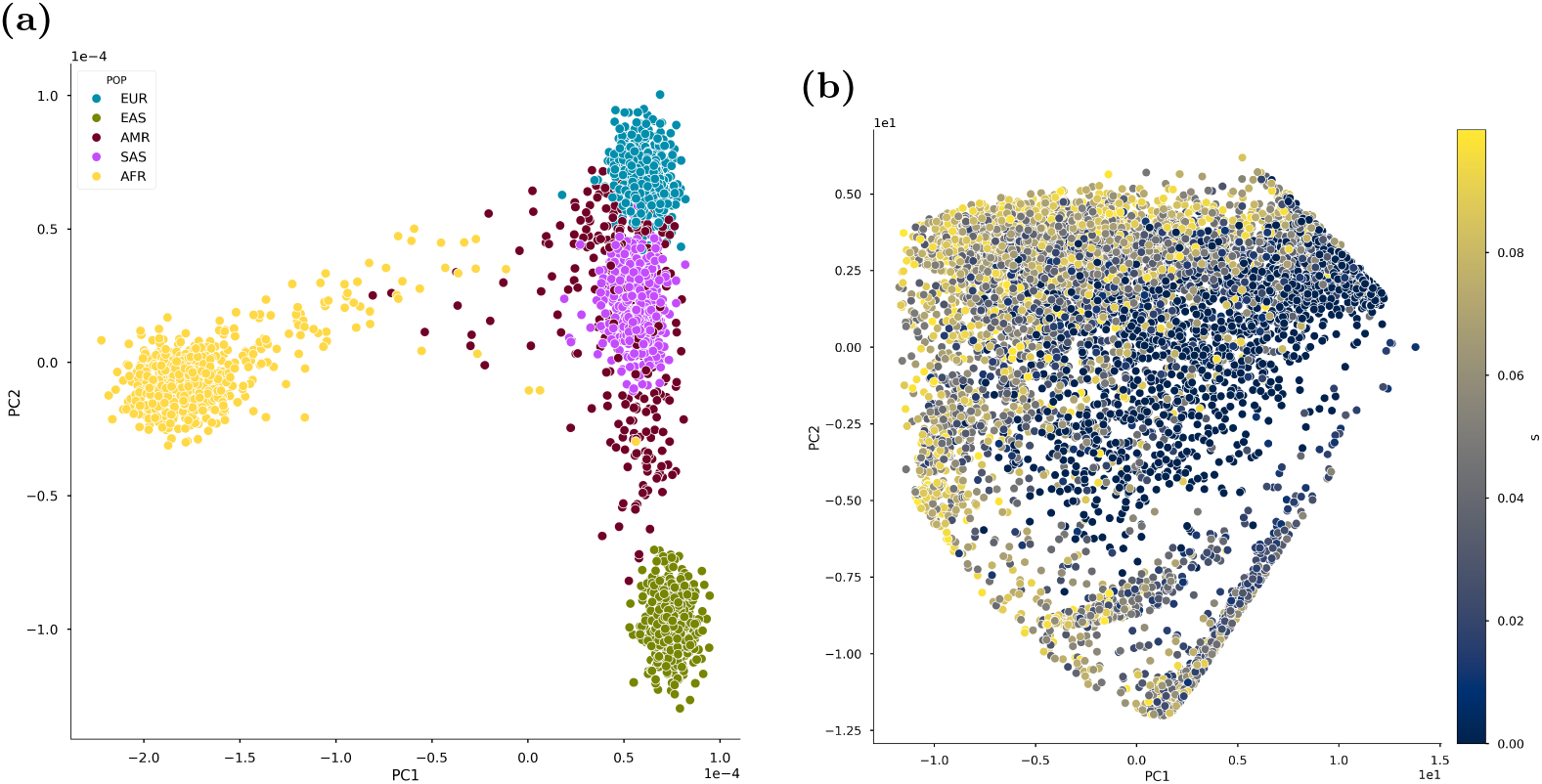
**(a)** PCA reduction of region-averaged learned embeddings for chromosome 1 for all 1000 Genomes populations, with colors indicating superpopulations (continent-level variation). Points represent chromosome 1 embeddings for diploid individuals. Population codes: EUR (Europe), EAS (East Asia), AMR (America), SAS (South Asia), AFR (Africa). **(b)** PCA reduction of window-level learned embeddings for simulated selection windows at a range of selection strengths. Points represent individual haplotype matrices.

Secondly, we aimed to identify whether the pre-training objective aligned at all with the task of selection detection. To do this, we calculated window-level embeddings for our selection simulations, which span a selection coefficient of 0-0.1 (Fig 3b). We observe that there is a clear gradient between low and high selection strengths. While not definitive, this indicates that the model’s pre-training task is partially aligned with the selection classification objective.

### Fine-tuned Popformer can accurately detect selection

We trained all of the supervised models on our human European-inferred (CEU) demographic simulations (for details on the simulations, other models, and training see Methods). In this study, we only aimed to detect positive selection, but training data encompassing negative or balancing selection could also be used. Each model classifies these simulated haplotype windows as either containing a selected-for mutation or not. We additionally include shouldering windows from selection simulations as negatives, aiming to improve localization power.

We report AUC and PR curves on a held-out test set of European simulations, and find that our method, in all training configurations, outperforms other models (fine-tuned, AUROC=0.95, AP=0.95). Fig 4). We additionally stratify the metrics by selection strength and by whether windows are shoulders (to a window containing a selected allele) and find that stronger selection is easier to detect, except for when the window is in a shoulder region of a stronger selected allele. These shoulders of strong selection are likely a strong confounding signal (S3 Fig).

**Fig 4.**
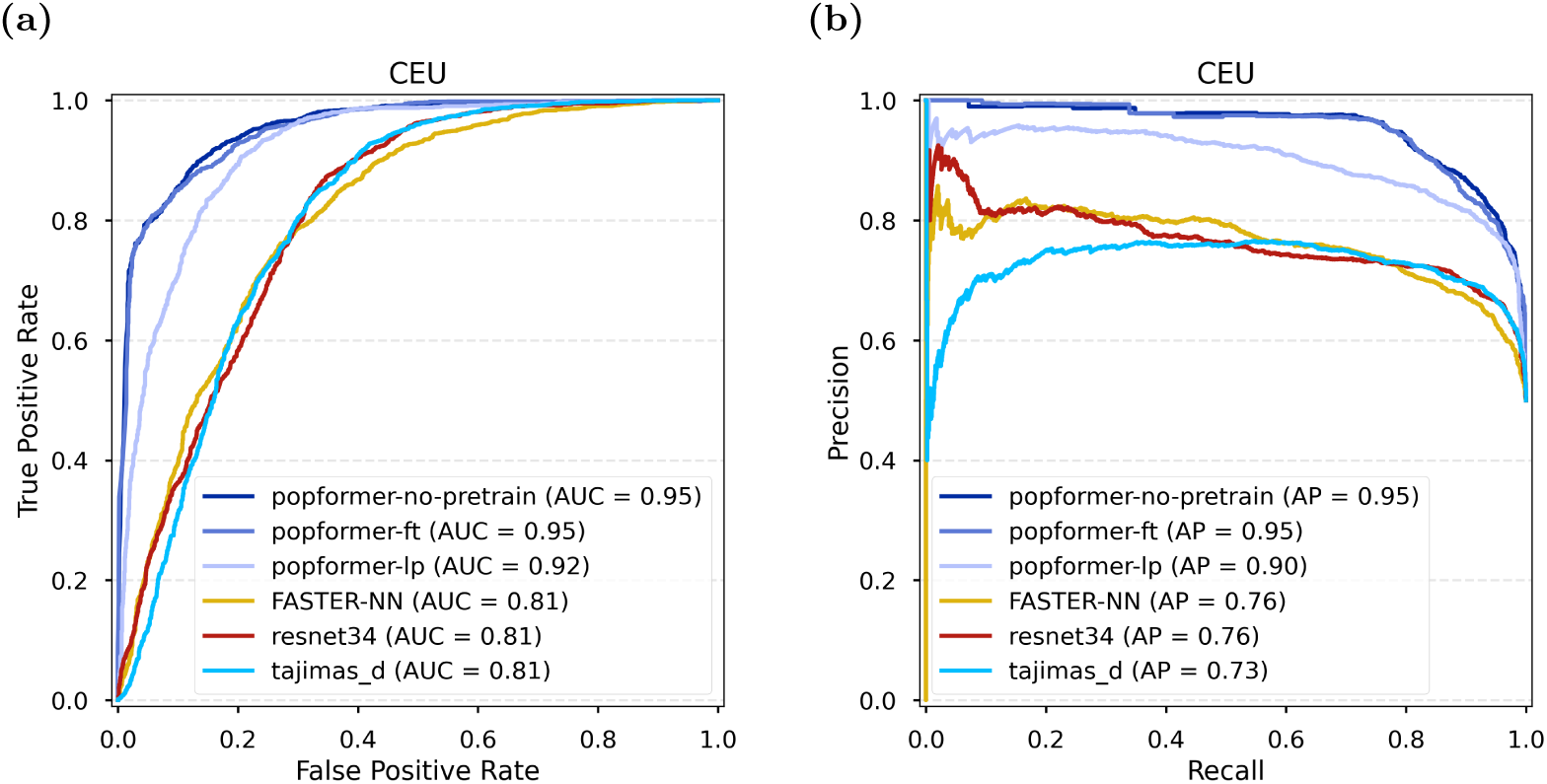
**(a)** ROC curves for our Popformer models, FASTER-NN CNN model, resnet34 CNN, and a summary-statistic (Tajima’s D) score, evaluated on a held-out test set of CEU-inferred demographic simulations. **(b)** Precision-recall curves for the same set of models and dataset.

While these results demonstrate the effectiveness of our model’s architecture at classifying selection from in-distribution simulations, the true benefits of the pre-training strategy are in the method’s ability to generalize to out-of-distribution (OOD) data. We evaluated all models on simulations from East Asian-inferred (CHB) and African-inferred (YRI) demographies, to evaluate models on their performance under demographic misspecification. We find that all methods perform roughly consistently with their performance on the in-distribution test set (Fig 5a, b).

**Fig 5.**
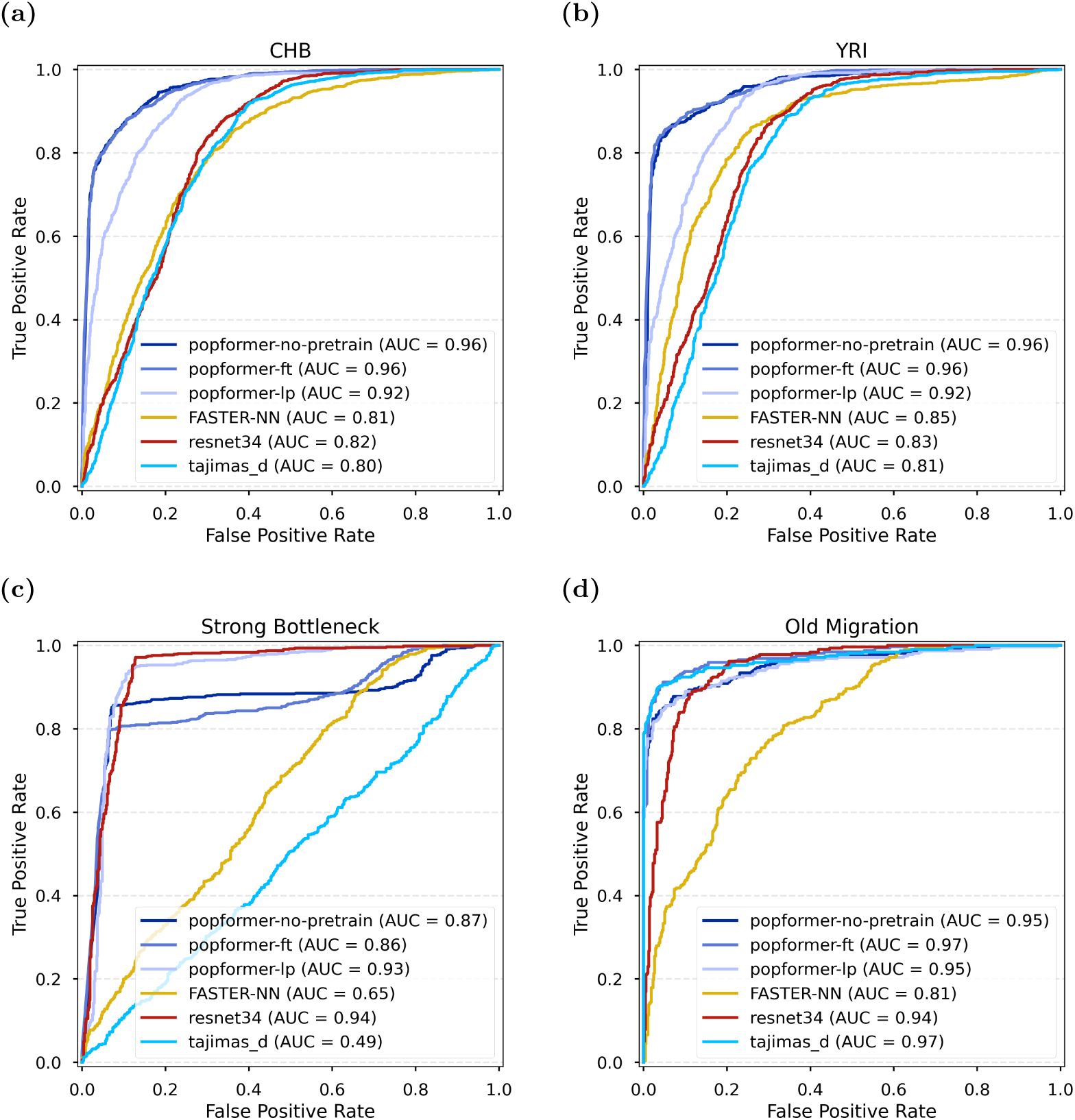
ROC curves for all models, evaluated on various simulated selection test sets. **(a)** CHB-inferred demographies. **(b)** YRI-inferred demographies. **(c)** Extremely strong bottleneck scenario. **(d)** Old migration demographic scenario.

We also test the models on dramatic misspecification, including an extremely strong bottleneck and an old migration scenario, which are signals confounding for selection [4, 36]. We report ROC curves and find that our model outperforms other models greatly in the difficult strong bottleneck scenario (Fig 5c, d). In the old migration scenario, we find that all methods perform extremely well, suggesting that the demographic scenario tested is not challenging. Despite that, fine-tuned Popformer still performs slightly better than the other deep-learning models and comparably with Tajima’s D.

Overall, Popformer models outperform the other methods we tested. Within the Popformer family, we find that the linear probe model (popformer-lp, frozen encoder) consistently performs worse than models which allow the encoder to update. This suggests that the pre-training task does not align perfectly with the selection classification task. Comparing the fine-tuned model (popformer-ft) with the ablation of the pre-training (popformer-no-pretrain), we do not find evidence suggesting that the pre-training helps with performance on simulated datasets. Instead, it might only benefit when applied to real data. We hypothesized that pre-training might benefit in a limited-data regime, but we also do not find this to be true (S4 Fig). The models order by performance does not change when varying the training dataset size.

### Validation on known selected regions

Though we display promising results on all simulated tests, ultimately, real evolutionary effects encompass a wider range of variation then we can simulate. Traditionally, methods are validated by inferring selection on a list of known or potentially novel sites, and examining whether predictions match the literature. We follow this tradition and analyze the commonly used LCT/MCM6 region (S6 Fig), which has a strong signature of positive selection in European populations [37].

However, this approach is prone to bias, in the form of cherry-picking a set of regions that a method identifies well, and does not provide any information on the false positive rate of a method. This approach is also uninformative on how powerful a model is, as any signals which are already well known are likely strong. We propose that selection detection methods, which are potentially most well suited for discovery (hypothesis generation), should focus heavily on controlling the false positive rate. The major difficulty with this is the fact that it is impossible to know the full ground truth for selection in the human genome.

Here, we introduce two approaches to validating selection detection methods on real data which can be used to compare methods on both power and false positive rate. First, new ancient DNA data and methodology has allowed for selection inference using real allele frequency trajectories [38]. Using these results, we gather a list of putatively neutral regions (*X <* 2) from Akbari *et al.* [29] to use as ‘true negatives’. By using the regions from Grossman *et al.* [28] as ‘true positives’, we can generate a pseudo-ROC-curve with the goal of comparing how well methods can differentiate between selected regions and putatively negative regions.

For most populations, which lack sufficient ancient DNA, we can instead compare [28]’s true positives with the fraction of the genome predicted to be under selection, under the assumption that a majority of the genome is neutrally evolving. Both of these approaches can be interpreted as analogous to a standard ROC curve, allowing for simple comparisons. These approaches are also independent of binary prediction threshold, which is important for deep learning methods, whose outputs are rarely calibrated.

Using these approaches, we find that the deep learning methods’ strong performances on simulated data do not translate well to performance on real data (Fig 6). Notably, Tajima’s D is the method which typically aligns best with the list of selection regions from [28], with the deep learning methods falling behind. Of the deep learning methods, however, Popformer generally recovers the known regions the best. However, our Popformer method outperforms all other methods, including Tajima’s D, at recovering known positive regions in the Yoruban (YRI) population. This result only holds in the fine-tuned Popformer training recipe (S5 Fig), not in the pre-training ablation, suggesting that real pre-training does provide an improvement when applying downstream models to real genomes. We likely only observe this benefit in YRI due to being the population most different from the European population used during training.

**Fig 6.**
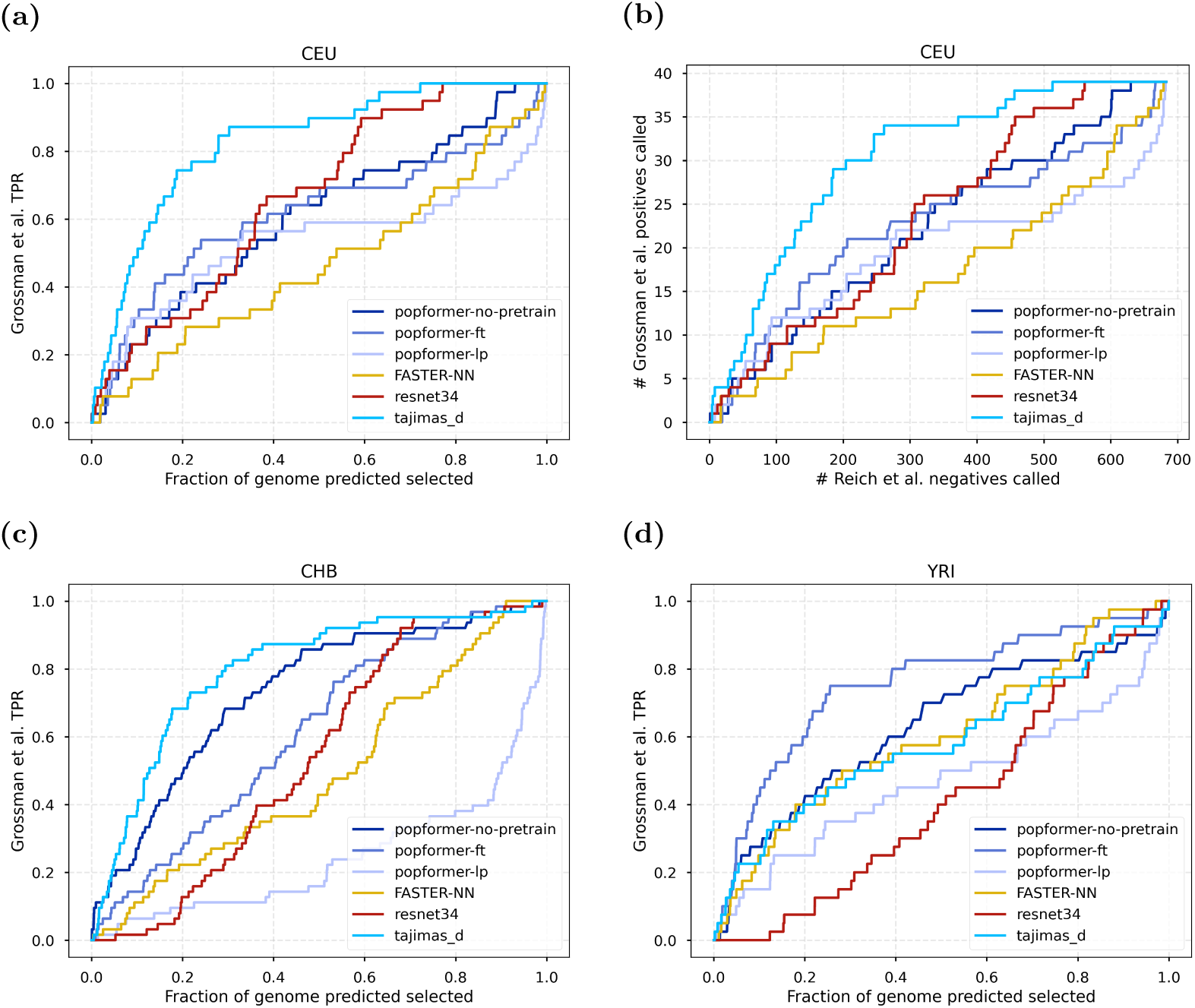
**(a)** All models, evaluated on recovery of Grossman *et al.* [28] known selection regions in the CEU population versus fraction of the genome predicted to be under selection. **(b)** Models evaluated on recovery of Grossman *et al.* CEU regions versus false discovery of Reich *et al.* negative regions. **(c)** Same as (a), but on the CHB population **(d)** Same as (a), but on the YRI population.

These results are representative of the fact that the selection signatures can manifest differently under different demographic models, and suggest that deep learning methods may be learning specific, not general, signatures. To begin to understand which, we examine correlations between Popformer-ft’s model scores and summary statistics, both for simulated (S7 Fig) and real (S8 Fig) datasets. There are some moderately sized correlations, which are expected (Tajima’s D = −0.68, # singletons = 0.56; Spearman correlation).

We also find evidence to support that using non-reported-positive regions as neutral regions is valid. Using inferred neutral alleles from ancient DNA as true negatives strikingly resembles using the fraction of the genome called (Figure 6a, b). Thus, if one is confident that their list of selected regions are true, using the rest of the genome as a neutral baseline in a test for recovery is likely to be informative. These analyses do not take into account potentially undiscovered selected regions in the genome – but, given that the reported regions are likely to be strong signals, detection methods should be able to predict them at higher probability than weaker and neutral signals.

## Discussion

Learning signatures of natural selection using supervised machine learning can improve selection detection power over traditional summary-statistic based methods [14, 16, 17, 36]. However, all simulation-based methods suffer when the simulations are poorly matched to the real data. In contrast, our method uses a modern paradigm for self-supervised learning in order to first learn important features of variation in real genomic data before fine-tuning on a particular supervised inference task. In addition, Popformer’s transformer-based architecture allows for more complex learned representations of genomic windows, including both relationships between SNPs and between haplotypes. Together, the self-supervised pre-training and the novel architecture enable Popformer to capture informative signals of selection. We demonstrate a higher windowed classification performance against two other deep-learning methods and a summary statistic score method.

Despite using self-supervised pre-training on real data, Popformer is still reliant on simulations to provide a training dataset for selection. Any such supervised method will have inherent data biases due to the necessary nature of selecting prior distributions on demographic and evolutionary simulation parameters. We aimed to mitigate this in our simulations by using a wide prior for our training simulations that encompasses a variety of possible human-like evolutionary histories. However, the main idea behind pre-training on real data is aimed to tackle this problem. Therefore, we tested on several human-like simulation sets and dramatically different simulation sets and found that Popformer achieves a level of robustness to out-of-distribution data superior to the other tested deep-learning methods and summary statistics. Ultimately, we are most interested in applying Popformer to real genomic datasets. We demonstrated that our method can modestly recover previously reported results of selection in three distinct human populations, but generalizes well to some populations (YRI) despite poorly matched training data (CEU-inferred). We believe that empirical validation of the results of a simulation-based selection method is highly important to assessing its usefulness and propose our validation strategy as one way to achieve this.

We believe that the Popformer architecture and self-supervised training strategy are an important contribution to the field of deep-learning selection inference. There do however remain many avenues for potential developments. Many approaches to unsupervised and self-supervised pre-training have been developed, including autoencoders and contrastive learning [39, 40]. As our findings on masked pre-training are not definitive, these approaches might be key to developing higher quality latent representations of population genomics data. Regarding genomic data and its representation, an input scheme where additional information (say, nucleotide variation tokens like ‘*A > T*’ [41] or embeddings from a genomic language model [42]) might allow for a complex transformer architecture like Popformer to better capture patterns of variation. Selection detection additionally depends on context, up to an extent. Efficient linear attention implementations, of which there are numerous, might allow for longer windows and potentially improved performance [43, 44]. Another future direction could be better quantification of uncertainty, which could be accomplished through recent advances in neural posterior estimation (NPE) [45, 46].

While used here only in the context of selection classification, the idea of learning signatures of variation in real data could easily be applied to other population-level (say, by averaging across windows) and window-level inference tasks in population genetics, including inference of demographic parameters, recombination hotspots, and mutation rates. These examples would be plug-and-play given a set of simulated training data and a pre-trained model, and would likely demonstrate the same robustness to mismatched data generation and assumptions as we found for selection. In addition, we believe that the SNP-level and haplotype-level attention architecture we introduce with Popformer would lend itself well to SNP-level population genetics tasks, including local ancestry inference or archaic introgression detection. Notably, Popformer could feasibly infer selection directly at a per-SNP level rather than a windowed level, allowing for greatly increased resolution. In fact, the pre-training masked language model objective we used might be more well suited to the granularity of these tasks. We expect that continual advancements in simulations and advances in genomic data quality and scale will reveal new applications and benefits of self-supervised deep-learning methods in population genetics.

## Materials and methods

### Details of the Popformer architecture

Popformer, in a similar manner to other raw-data deep-learning methods, operates on matrices **x** ∈ {0, 1}*^n^*^×^*^S^* of *n* haplotype rows by *S* single nucleotide polymorphism (SNP) columns derived from genomic windows. Only biallelic SNPs are included. These matrices are computed in the form of binary matrices in which 0 represents the more common (major) allele and 1 represents the less common (minor) allele. We also include a vector *d* ∈ N*^S^* encoding distances between SNPs, which allows the model to incorporate positional information in its predictions. Unlike most CNN methods, the transformer architecture can handle both a variable number of haplotype rows and SNP columns without additional padding. This renders Popformer more efficient when performing inference on populations that have a different number of individuals.

An overview of the architecture can be found in Fig 1. We adopted the model architecture from the axial attention transformer architecture, originally developed for image learning and later for protein sequence learning by MSA Transformer [30, 31]. Attention is a widely-used method that allows each element in an input sequence to be dynamically weighted (attended to) based on every other element in the sequence. For example, text tokens in a sentence can each be weighted based on the context from the full sentence. 2D axial attention, originally designed for image methods, extends the standard mechanism of attention, which operates on 1-dimensional sequences, to operate consecutively across multiple dimensions [30]. Column-wise attention (in our case representing structure across haplotypes) is applied for every SNP with complexity *O*(*Sn*^2^). We use tied row attention, introduced by [31], to share the row-wise self attention maps between every haplotype in the matrix. This forces the model to learn one pattern of SNP dependency, which maps directly between different haplotypes. It also reduces memory usage from *O*(*S*^2^*n*) to *O*(*S*^2^). Following most transformer architectures, we include residual connections around the attention layers [47]. In all, each axial attention block consists of a row-attention block, a column attention block, and a feedforward dense network, allowing us to aggregate information about variant and population structure.

In the standard self-attention implementation, a positional embedding of a number of forms can be used to break standard attention’s positional-invariance. These forms all assume that ‘distances’ between consecutive tokens are equal. Unlike protein and image sequences, adjacent columns of SNPs in the input matrices do not represent consecutive positions within the original genome. The distances between variants provide SNP density information otherwise lost in a pure haplotype matrix. In order to encode the meaningful patterns of inter-SNP distances, we include an altered form of the T5 language model’s learned positional embedding approach [48]. Inter-SNP genomic distances (in base pairs) are first binned into a fixed number of linearly-spaced distance buckets. An embedding is learned for each bucket separately for each row-attention head and for each layer. The embedding directly modifies the attention matrix in the form of an added bias. We adapt T5’s 32 learned positional embeddings, using 31 linearly-spaced discrete distance buckets from 0 to 50k base pairs, with a final 32nd bucket reserved for any larger distances.

### Pre-training on real genomes

Our pre-trained model (Popformer-base) is trained with an analogue of the masked language modeling objective. Given masked positions *M* ⊆ {1 … *n*} × {1 … *S*} using which the masked input **x̃** are created, we aim to find model parameters *θ* which minimize the binary cross-entropy loss:

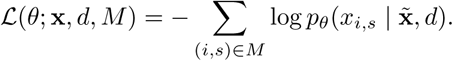

The model outputs logits for the major and minor alleles for every position, while the loss is computed only for masked tokens. We do not mask *d*, the distance vector. In our case, this masking is applied uniformly across all SNPs (S1 Fig). We use a high masking rate (75%) to prevent models from learning simple extrapolation from neighboring haplotypes or alleles, as in [49]. We experimented with additionally masking columns of SNPs, blocking the model from using context from different haplotypes, but find that this degrades downstream model performance. We additionally tried to weight rare variants by oversampling minor alleles (1s) during masking. This resulted in poor performance on validation sets as the model learned from an incorrect distribution of major/minor alleles.

The training data for Popformer’s pre-training contained 260,000 haplotype windows, sampled uniformly from all 26 populations of the 1000 Genomes phase 3 release [32]. A random population and start position along the genome are chosen. We find the nearest SNP to the start position, and perform basic quality control by reselecting a new window if less than 50% of the bases in the window are in the 1000 Genomes phase 3 strict accessibility mask. For each window, we select a SNP window size uniformly between 32 and 512 (only including biallelic SNPs) and a number of haplotypes uniformly between 32 and 128. Thus, haplotypes within a window are guaranteed to be from the same population but their order and number is randomly selected.

For all Popformer results, we used a model with four hidden layers, with each layer containing four self-attention heads each for row- and column-attention (34M parameters). This model is relatively smaller than similar models used in the vision, protein, and language domains, as we both have less complex data representations and less training data. Cross-entropy loss was used during both pre-training (at the per-SNP level) and during fine-tuning (at the window level). The Adam optimizer was used with a linear learning rate decay and a warmup period of 0.1 epochs. This pre-trained model (Popformer-base) is used as the base model for the downstream selection task by adding a simple linear classification head. We tested two downstream training strategies: full fine-tuning, in which all the weights of the model (including the transformer encoder) are allowed to update, or training only the linear head (with the encoder frozen, referred to as linear probing). We additionally test a model with ablated pre-training, in which we trained a randomly initialized encoder and linear head from scratch on the selection detection task.

In all, the pre-training took 6 hours to complete on a B200 GPU with 128 GB of memory. Fine-tuning took 1 hour to complete on a single RTX 5000 GPU (32 GB). Inference over 49,000 64-haplotype 50kbp windows (genome length) took 7 minutes to complete on the same RTX 5000 GPU.

### Selection simulations

We used the `SLiM` simulation framework through the high-level `stdpopsim` library to generate samples for training and testing [22, 50, 51]. To cover a wide range of demographic parameters, we use demographic parameters randomly sampled from inferred demographies for CEU, CHB, and YRI populations using `pg-gan`, a GAN-based approach to demographic inference [52]. We filter sets of demographic parameters which have growth rate *<* 0.01 for ease of simulation. A summary of demographies used is in S1 Table. We sample recombination rate uniformly at random from the HapMap recombination map and sample mutation rate randomly in *µ* ~ *U* (1.29 × 10^−8^*/*2, 1.29 × 10^−8^ × 2) [53]. We simulated a total of 5,000 neutrally evolving regions and 5,000 regions with a centered beneficial allele, all of length 250kbp. We randomly varied the strength of selection in *s* ~ *U* (0.01, 0.1), and chose an onset time such that the allele would roughly reach fixation by the end of the simulation using Equation A17 of [54].

For each neutrally evolving sample, we randomly choose 10 sub-windows, all of which are labeled as neutral. For each selected sample, we randomly choose 10 sub-windows not containing the selected allele to be labeled as neutral (shoulders). We choose 20 sub-windows containing the selected allele to be labeled as positive. In all, this gives us 100,000 total neutral windows and 100,000 total positive windows. These are split first into populations, and for each population into an 80/20 train/test split. In all, these human-inferred simulations took 5 hours on 20 CPUs and storage of raw genotypes took 10GB on disk.

In addition, we simulated two potentially confounding demographic scenarios to test the ability of each model to generalize to non-training examples - 1) an extremely strong bottleneck with severity 0.005 following [36] and 2) a scenario with very old migration. The old migration scenario involves a population split 2,000 generations ago, with the two derived populations admixing (in 80/20 proportions) 1,000 generations ago. These simulations were only used for testing.

### Selection methods

We found it extremely important to implement fair model comparisons. Most machine learning selection classifiers are not designed to be used out-of-the-box, but rather trained from scratch on a target demographic model. Thus, we designed an evaluation harness for training and evaluating window-based classification methods that facilitates making fair model comparisons as we did in this work. This harness was also designed to be extensible, allowing for additional models to be added with minimal effort, and is released along with the models at https://popformer.leonzong.com.

We compared our method with two deep-learning methods and Tajima’s D. A widely used architecture for selection detection deep-learning models are convolutional neural networks (CNNs). CNNs are naturally suited to the haplotype input data, both due to the input shape and due to the fact that learning convolutional filters aggregates local patterns of diversity in an appropriate way for discovering local signatures of sweeps.

FASTER-NN is a 1D-CNN that takes as input 2 1D vectors - minor allele frequencies and distances [36]. We use the same training recipe as was used in the original FASTER-NN method, except using our training simulations. We additionally compared against a ResNet architecture, which are CNNs commonly used for images. For this model, we adopt the preprocessing steps from [16, 55] in which we sort haplotypes based on genetic similarity. This model was trained with a batch size of 16 and learning rate of 1e-4 for around 20 epochs, at which point validation loss stopped improving.

## Supporting information

**S1 Fig. An example masked window.** From top to bottom: 75% uniformly masked input, predicted output, and ground truth labels. These reflect Popformer’s self-supervised pre-training process. Loss is computed between the predicted and ground truth for masked positions and used to update the model’s weights.

**S2 Fig. Example genotype imputation test window.** The first 64 haplotypes are in the “reference” panel and are unmasked, and the next 64 haplotypes are the “target” panel and are randomly masked.

**S3 Fig. Stratified performance of all models.** ROC curves for models on held-out CEU-inferred test set, stratified by selection strength (columns) and by shoulders (rows, included shoulder regions or excluded).

**S4 Fig. Performance of models at varying training data amounts.** Average precision (AUPRC, equivalent to AP) of all models at varying percentages of training dataset used. All models were trained to convergence.

**S5 Fig. Comparisons of Popformer training recipes.** [28] set of positives vs total genome predicted under selection for each of CEU, CHB, and YRI, only comparing different Popformer training recipes.

**S6 Fig. Windowed predictions around the LCT/MCM6 allele in the CEU population.** The highlighted region represents the selection region reported in [28]. Each prediction line is for a 50kbp window.

**S7 Fig. Correlations between Popformer scores and simple summary statistics in simulations.** Scatterplots of Popformer-ft scores and Tajima’s D, # of singletons, and number of SNPs in window (SNP density), on held-out CEU-inferred test set. Points represent windows and are colored by selection strength.

**S8 Fig. Correlations between Popformer scores and simple summary statistics.** Scatterplots of Popformer-ft scores and Tajima’s D, # of singletons, and number of SNPs in region (SNP density), on all real-CEU-genome windows.

**S1 Table Parameter ranges used for per-population inferred human demographic simulation.** 20 total demographic models for each population were inferred, and only those with growth rate *<* 0.01 are used as possible demographies for our simulations.

## Acknowledgments

This work is funded by NIH grant R15HG011528. The content is solely the responsibility of the authors and does not necessarily represent the official views of the National Institutes of Health.

