## Supplementary Material for "Popformer: Learning general signatures of positive selection with a self-supervised transformer"

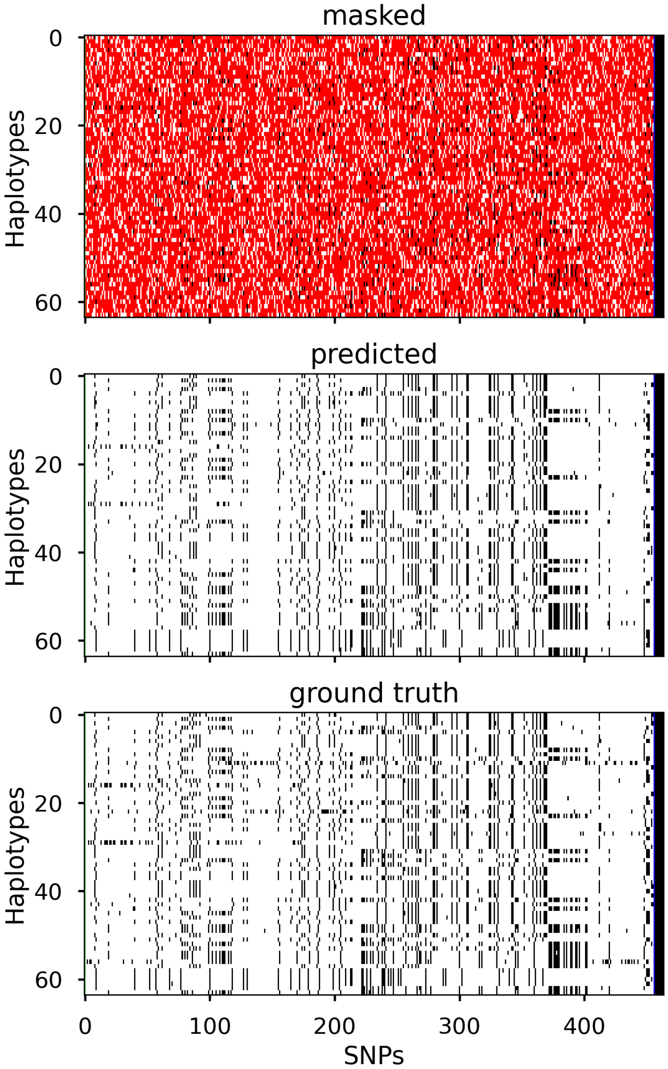


**S1 Fig. An example masked window.** From top to bottom: 75% uniformly masked input, predicted output, and ground truth labels. These reflect Popformer's self-supervised pre-training process. Loss is computed between the predicted and ground truth for masked positions and used to update the model's weights.

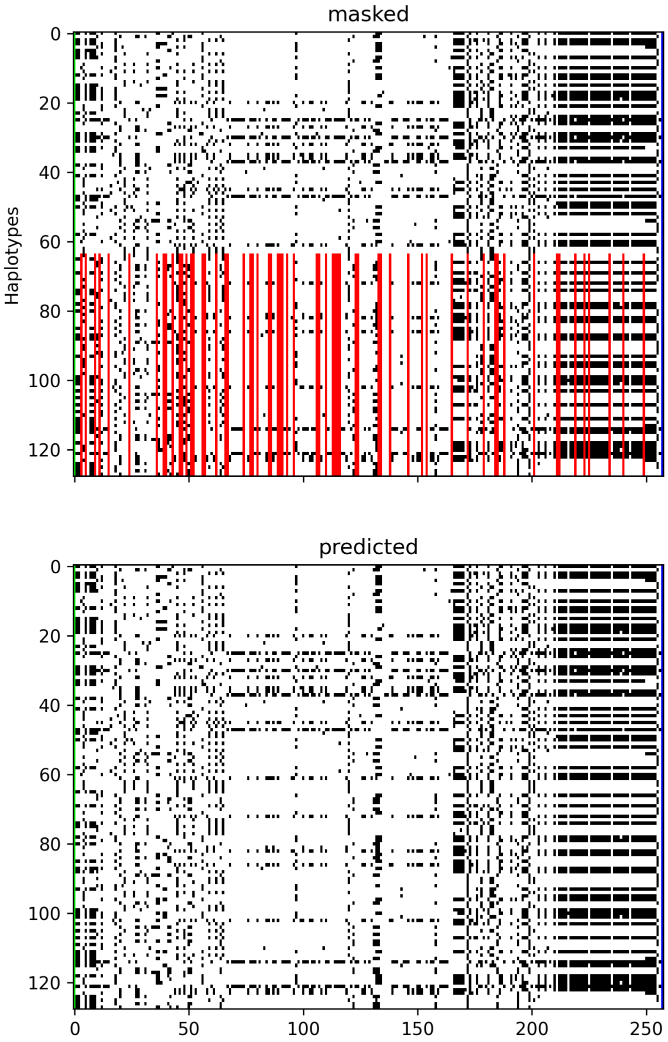


**S2 Fig. Example genotype imputation test window.** The first 64 haplotypes are in the "reference" panel and are unmasked, and the next 64 haplotypes are the "target" panel and are randomly masked.

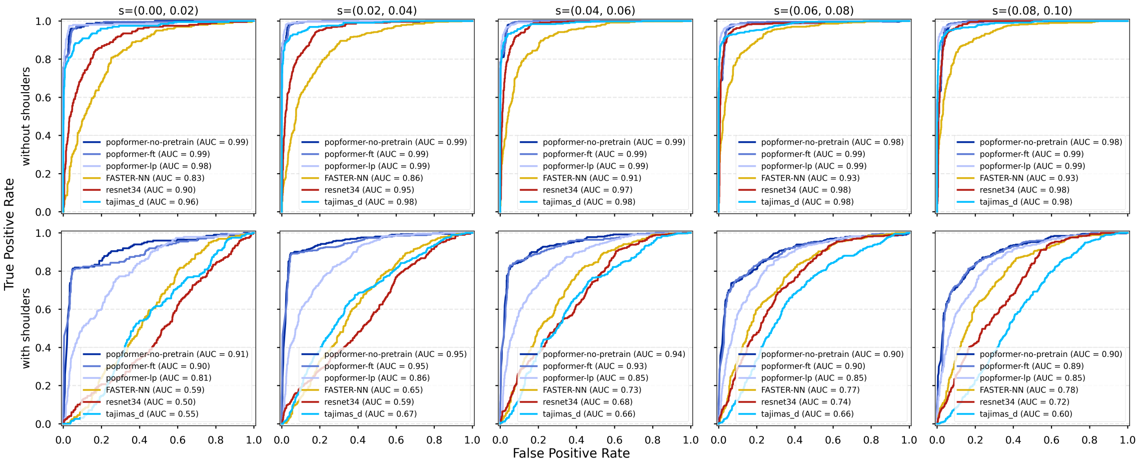


**S3 Fig. Stratified performance of all models.** ROC curves for models on held-out CEU-inferred test set, stratified by selection strength (columns) and by shoulders (rows, included shoulder regions or excluded).

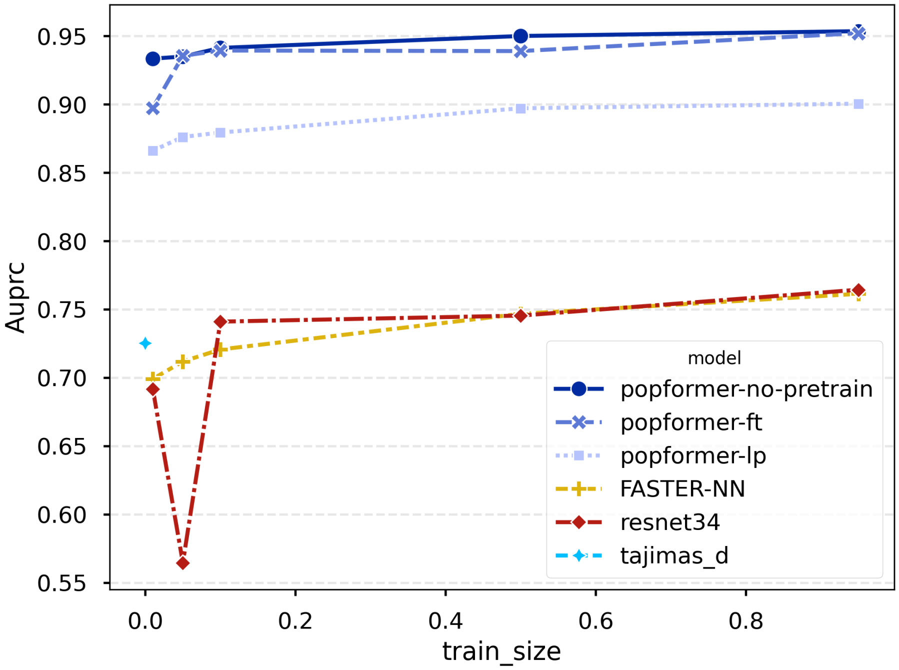


**S4 Fig. Performance of models at varying training data amounts.** Average precision (AUPRC, equivalent to AP) of all models at varying percentages of training dataset used. All models were trained to convergence.

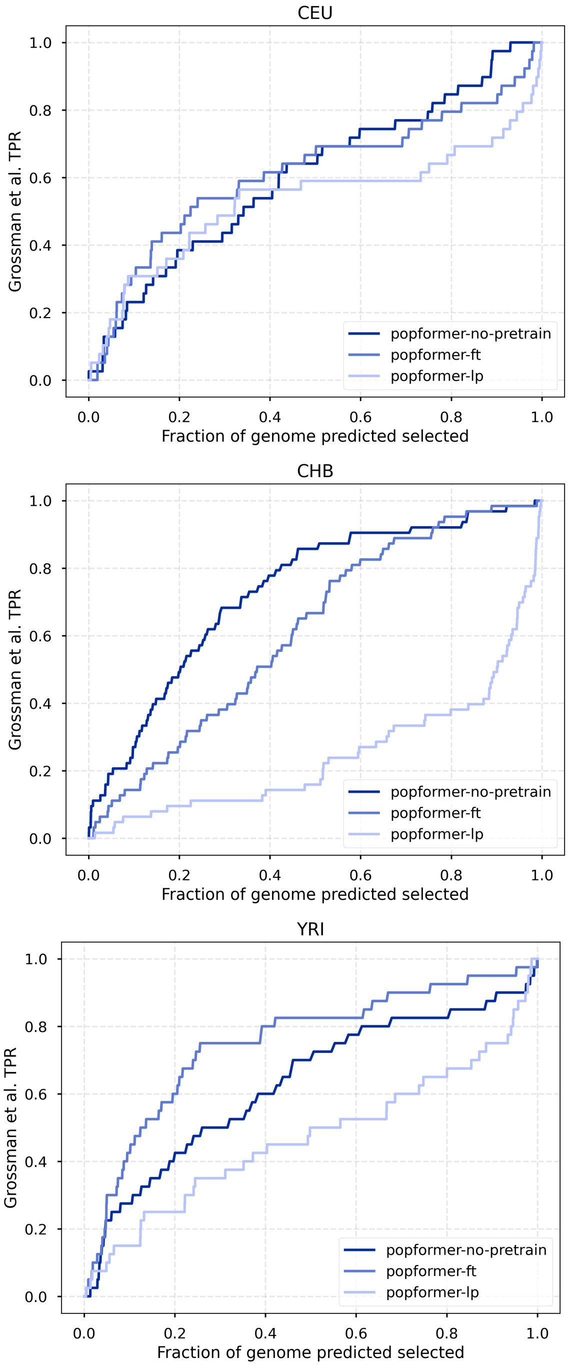


**S5 Fig. Comparisons of Popformer training recipes.** Grossman et al. set of positives vs total genome predicted under selection for each of CEU, CHB, and YRI, only comparing different Popformer training recipes.

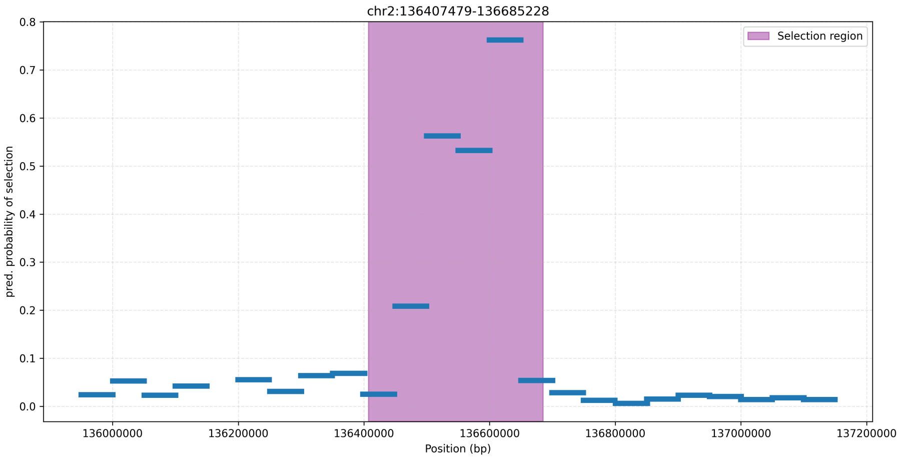


**S6 Fig. Windowed predictions around the LCT/MCM6 allele in the CEU population.** The highlighted region represents the selection region reported in Grossman et al. Each prediction line is for a 50kbp window.

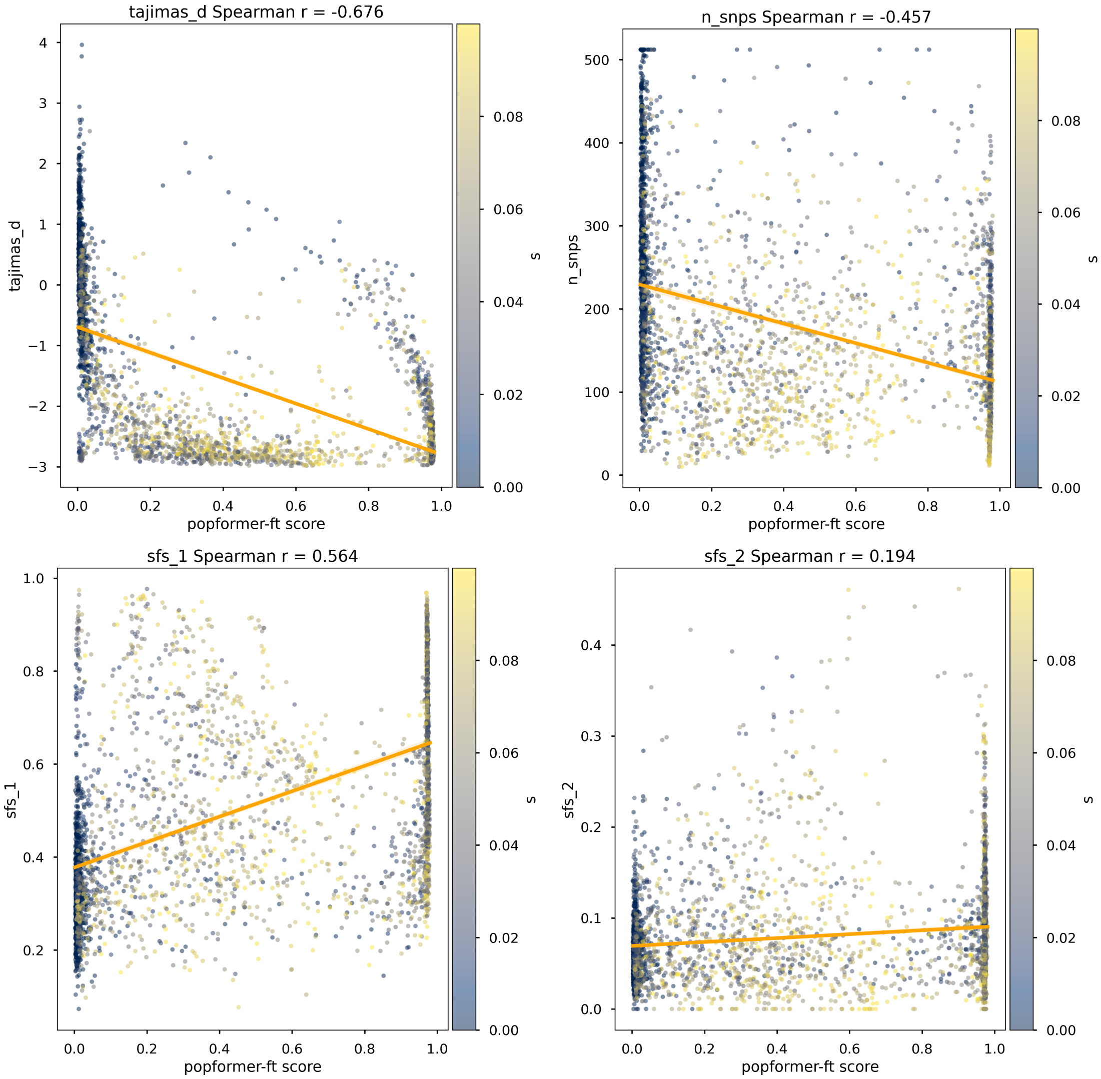


**S7 Fig. Correlations between Popformer scores and simple summary statistics in simulations.** Scatterplots of Popformer-ft scores and Tajima's D, number of SNPs in window (SNP density), # of singletons, and # of doubletons on held-out CEU-inferred test set. Points represent windows and are colored by selection strength.

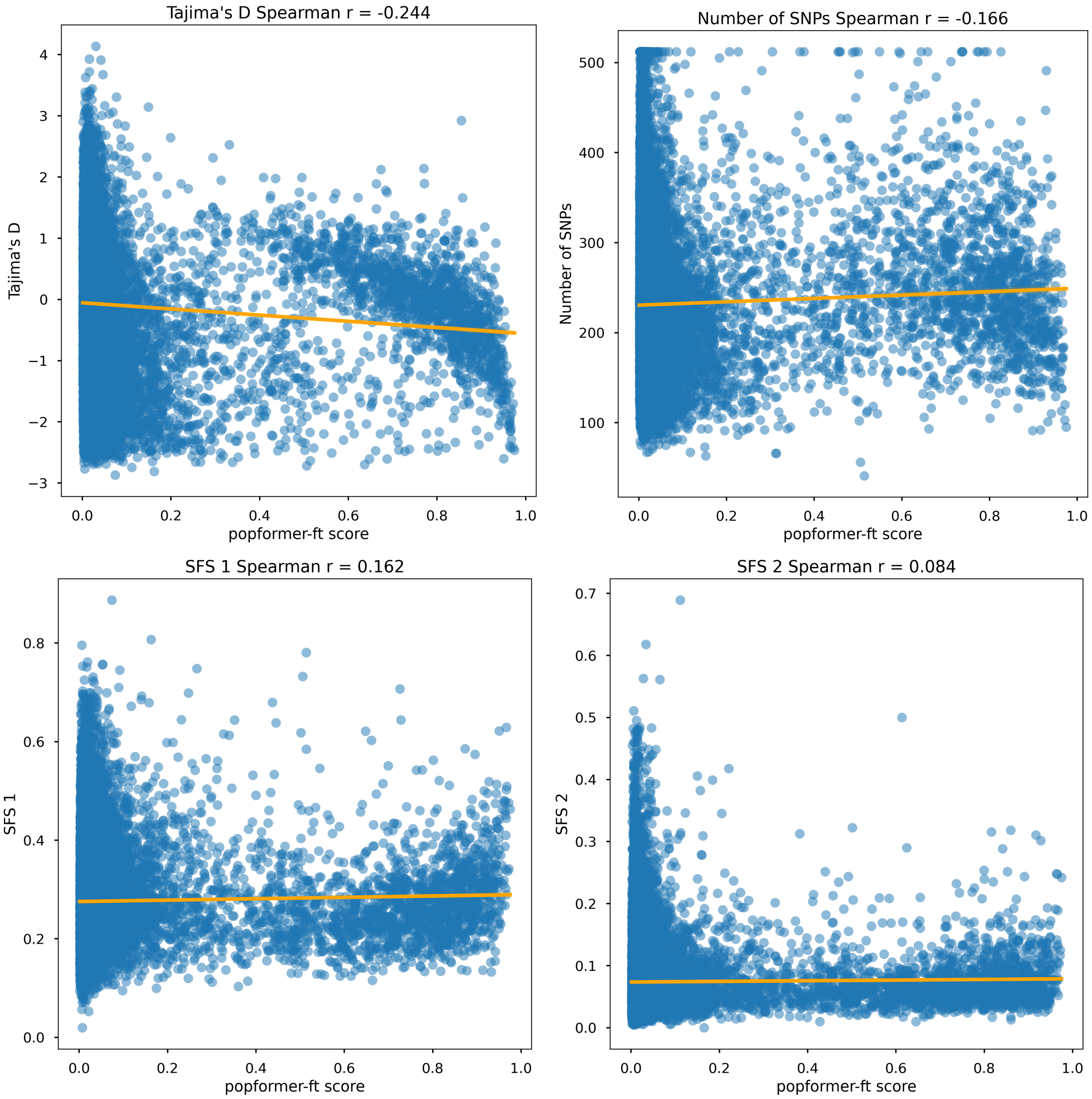


**S8 Fig. Correlations between Popformer scores and simple summary statistics.** Scatterplots of Popformer-ft scores and Tajima's D, number of SNPs in window (SNP density), # of singletons, and # of doubletons on all real-CEU-genome windows.

| pop | N1 | N2 | T1 | T2 | growth |
|---|---|---|---|---|---|
| CEU | 24034 | 3110 | 3261 | 1349 | 0.0032 |
| CEU | 21775 | 3216 | 3270 | 938 | 0.0064 |
| CEU | 18344 | 7044 | 3619 | 784 | 0.0030 |
| CEU | 21370 | 4001 | 3239 | 871 | 0.0065 |
| CEU | 18242 | 8019 | 3390 | 559 | 0.0093 |
| CHB | 24694 | 8298 | 3889 | 876 | 0.0020 |
| CHB | 25107 | 5075 | 4863 | 1354 | 0.0029 |
| CHB | 23106 | 4270 | 4042 | 694 | 0.0098 |
| CHB | 14941 | 4285 | 1967 | 1414 | 0.0017 |
| YRI | 23049 | 22564 | 3620 | 516 | 0.0040 |
| YRI | 21880 | 28129 | 4024 | 511 | 0.0024 |
| YRI | 26962 | 12884 | 4724 | 1043 | 0.0076 |

**S1 Table. Parameter ranges used for per-population inferred human demographic simulation.** 20 total demographic models for each population were inferred, and only those with growth rate $< 0.01$ are used as possible demographies for our simulations.
